## Supplemental Information for "Controllability Governs the Balance Between Pavlovian and Instrumental Action Selection"

**Derivation of Bayesian models**

The generative model of an uncontrollable environment assumed by the Pavlovian system is given by:

$$\theta_{s} \sim Beta\left( \theta_{0}\frac{\eta_{0}}{2}, \left( 1-\theta_{0} \right)\frac{\eta_{0}}{2} \right)$$

$$r|s \sim Bernouli(\theta_{s})$$

where $\theta_{s}$ is the reward probability for stimulus *s*, $\theta_{0}$ is the mean of the prior, $\eta_{0}$ is a parameter controlling the dispersion of the prior, and *r* is the reward outcome on a particular trial (all reward outcomes are assumed to be independently and identically distributed). The generative model of a controllable environment assumed by the instrumental system is essentially the same, except conditioned on action, *a*:

$$\theta_{sa} \sim Beta\left( \theta_{0}\frac{\eta_{0}}{2}, \left( 1-\theta_{0} \right)\frac{\eta_{0}}{2} \right)$$

$$r|s,a \sim Bernouli(\theta_{sa})$$

For both systems, the posterior conditional on the stimulus-action-reward history $\mathcal{D}$ is a Beta distribution with the same functional form as the prior, shown here for the Pavlovian system:

$$P\left( \theta_{s} | \mathcal{D,}m=uncontrollable \right)=Beta(\hat{\theta}_{s}\frac{\eta_{s}}{2},(1-\hat{\theta}_{s})\frac{\eta_{s}}{2})$$

$$\hat{\theta}_{s}\boldsymbol{=}\frac{{\eta_{0}+N}_{s}}{\eta_{s}}$$

$$\eta_{s}= \eta_{0}+T_{s}$$

where *m* indexes the assumed environment (uncontrollable for the Pavlovian system, controllable for the instrumental system), $N_{s}$ is the number of times stimulus *s* was paired with reward, and $T_{s}$ is the number of times stimulus *s* was presented. The equations for the instrumental system are identical except for conditioning on both stimuli and actions. Through simple algebraic manipulation, the update for the posterior mean $\hat{\theta}_{s}$ can be expressed as a recursive learning rule (Eq. 2 in the main text).

The posterior over generative models is given by:

$P\left( m=uncontrollable | \mathcal{D} \right)\propto P\left( \mathcal{D} | m=uncontrollable \right)P(m=uncontrollable)$.

The marginal likelihood is an integral over the latent parameters:

$$P\left( \mathcal{D} | m=uncontrollable \right)= \int P\left( \mathcal{D} | m=uncontrollable, \theta_{s} \right)P\left( \theta_{s} \right)d\theta_{s}= \frac{B(\hat{\theta}_{s}\frac{\eta_{s}}{2},(1-\hat{\theta}_{s})\frac{\eta_{s}}{2})}{B(\theta_{0}\frac{\eta_{0}}{2}, \left( 1-\theta_{0} \right)\frac{\eta_{0}}{2})}$$

where *B* denotes the beta function. The beta function can be expressed recursively:

$$B\left( x+1,y \right)=B\left( x,y \right)\frac{x}{x+y}$$

$$B\left( x,y+1 \right)=B\left( x,y \right)\frac{y}{x+y}$$

Applying these recursions to the log posterior odds over *m*, one obtains the update rule for *L* in the main text (Eq. 6).

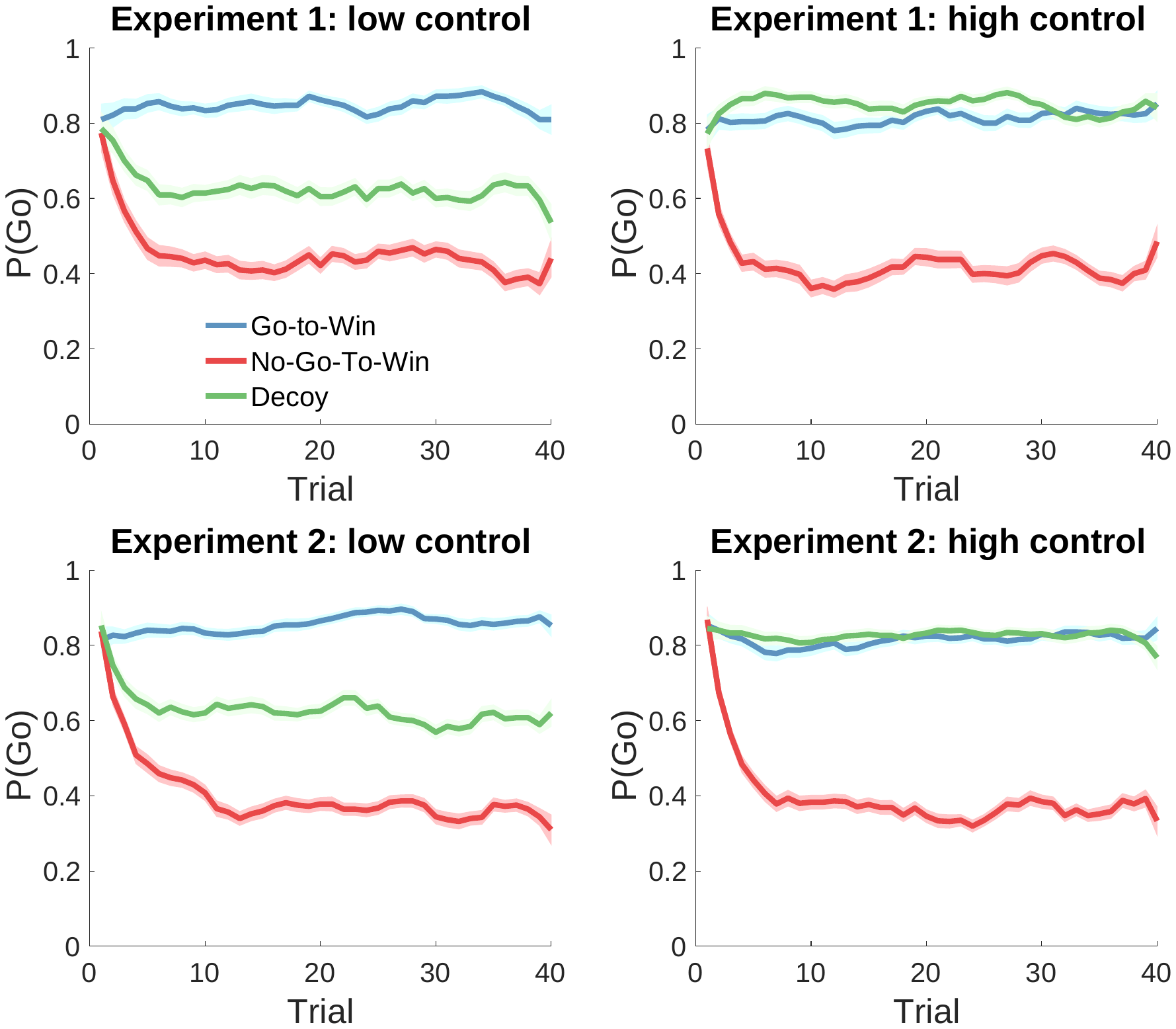

Figure S1. Probability of Go response across trials for each experimental condition and stimulus, smoothed with a 5-trial moving average. Error bars show standard error of the mean.
